# Metabarcoding replicate detection frequency tracks ddPCR copy number for cod and herring eDNA in ancient marine sediments

**DOI:** 10.64898/2026.07.03.736335

**Authors:** Elena Baños, Steen Wilhelm Knudsen, Kristine Bohmann, Luke E. Holman

## Abstract

1. Detecting environmental DNA (eDNA) from rare or low-abundance aquatic species remains a major challenge, particularly when it is highly degraded, present at low concentrations, and dominated by DNA from non-target taxa. These challenges are further amplified in sedimentary ancient DNA (sedaDNA) studies, where thousands of years can degrade eDNA further, making the detection and quantitative interpretation of weak biological signals difficult.
2. Metabarcoding is commonly used to produce high-throughput community-level data from eDNA but is inherently compositional and influenced by amplification biases. Nonetheless, metabarcoding read abundance or PCR replicate detection frequency are increasingly used as proxies for relative DNA concentration, but their quantitative interpretation has rarely been evaluated against independent measures of absolute DNA abundance.
3. We used droplet digital PCR (ddPCR) to quantify mitochondrial DNA from Atlantic cod (*Gadus morhua*) and Atlantic herring (*Clupea harengus*) in 136 ancient eDNA extracts from Icelandic marine sediment cores spanning the last three millennia. We compared ddPCR copy number estimates with metabarcoding (18S) derived relative abundance and detection frequency, and evaluated whether temporal DNA trends corresponded with proxy reconstructed sea surface temperature (SST) variability.
4. We found that ddPCR-measured fish sedaDNA abundance was positively correlated with the proportion of metabarcoding PCR replicates for both Atlantic cod and Atlantic herring. Moreover, temporal trends in Atlantic herring DNA abundance were consistent with proxy reconstructed SST variability, supporting the ecological relevance of the molecular signal.
5. Overall, our results show that ddPCR-derived DNA concentrations and metabarcoding PCR replicate detection frequency capture consistent patterns in low-abundance fish sedaDNA from marine sediments. The observed agreement between approaches supports the use of PCR replicate detection frequency as a semi-quantitative proxy for low-abundance sedaDNA.

## 1 Introduction

Biodiversity underpins ecosystem function and resilience, sustaining the ecosystem services on which human societies depend (Cardinale et al., 2012). Safeguarding these benefits requires an understanding of ecosystem change across time, and thus a long-term perspective is essential to evaluate species persistence, community turnover and to define meaningful conservation baselines (Bálint et al., 2018; Gillson et al., 2023). Reliable methods to detect species deep into the past are therefore critical for assessing biodiversity dynamics and anticipating responses to future environmental change or anthropogenic pressures (Lindahl et al., 2025). One approach is the extraction and analysis of ancient environmental DNA (eDNA) preserved in sedimentary cores, commonly referred to as sedimentary ancient DNA (sedaDNA), which can reliably reconstruct past biodiversity across millennial timescales (Capo et al., 2022; Nguyen et al., 2023).

A key challenge in many sedaDNA studies, particularly those targeting multicellular taxa, is that target DNA is often present at very low concentrations (Armbrecht et al., 2021; Giguet-Covex et al., 2019). This reflects limited DNA shedding relative to the total environmental DNA pool (Barnes & Turner, 2016), strong dilution by non-target DNA (Mauvisseau et al., 2022), and variable degradation rates across different taxa or sedimentary conditions (Brandão-Dias et al., 2025a; Sand et al., 2024). These processes can result in target DNA being rare, highly fragmented, and close to analytical detection limits, thereby complicating reliable detection and quantification. To address these challenges, multiple molecular approaches have been developed, including community-level methods such as metabarcoding (Holman et al., 2025a; Zimmermann et al., 2024), shotgun metagenomics (Zimmermann et al., 2023; Holman et al., 2025b), hybridisation-capture metagenomics (Armbrecht et al., 2021; Foster et al., 2024; Foster et al., 2026; Schulte et al., 2021), and targeted techniques such as quantitative PCR (qPCR) (Lopez et al., 2024) or droplet digital PCR (ddPCR) (Harðardóttir et al., 2024).

Metabarcoding is the most widely used molecular approach in sedaDNA studies (Capo et al., 2022; Holman et al., 2023). Through amplifying and sequencing homologous DNA regions, metabarcoding provides a high-throughput community characterisation from eDNA extracts (Armbrecht et al., 2020; Barrenechea et al., 2023; De Schepper et al., 2019). However, all metabarcoding datasets are affected by PCR amplification biases and are inherently compositional, limiting quantitative interpretation (Elbrecht & Leese, 2015; Kelly et al., 2016, 2019; Shaffer et al., 2025). These complexities are compounded in ancient eDNA, where DNA is typically highly fragmented and at low concentrations relative to abundant non-target environmental DNA (Orlando et al., 2015; Selway et al., 2022).

While there has been recent progress towards truly quantitative metabarcoding (McLaren et al., 2019; Shelton et al., 2023), most eDNA studies rely on indirect metrics to approximate DNA abundance (Chen & Ficetola, 2020; Herzschuh et al., 2025). The proportion of positive detections across multiple technical PCR replicates is increasingly used as a semi-quantitative proxy for target DNA concentration in sedaDNA studies (Alsos et al., 2024; Garcés-Pastor et al., 2022). This method assumes that PCR replicate detection frequency correlates with underlying DNA concentration, and has been shown to track ecological variation in sedaDNA records (Alsos et al., 2020; Ekram et al., 2025; Holman et al., 2025a), providing a semi-quantitative measure across many taxa (Alsos et al., 2024; Clarke et al., 2024; Garcés-Pastor et al., 2025; Zetter et al., 2025). However, its relationship to independently derived absolute measurements of DNA concentration remains unclear.

Droplet digital PCR (ddPCR) provides accurate quantification of target DNA molecules by partitioning the PCR into thousands of nanoliter-sized droplets before target detection (Hou et al., 2023; Sedlak & Jerome, 2013). It is highly sensitive, is less susceptible to PCR inhibition than other methods (Hamaguchi et al., 2018; Mejbel et al., 2021; Rački et al., 2014), and does not rely on DNA target standards for quantification, making it suitable for low concentration ancient DNA (Capo et al., 2021). Despite increasing use of ddPCR in modern eDNA research (Doi et al., 2015; Picard et al., 2022), it has been applied to sedaDNA in relatively few studies (e.g. Capo et al., 2021; De Schepper et al., 2019; Hamaguchi et al., 2018; Harðardóttir et al., 2024; Mejbel et al., 2021; Thomson-Laing et al., 2025). Moreover, the relative performance of ddPCR and metabarcoding for quantifying low-abundance species in ancient sedimentary records remains poorly understood, largely because available evidence is dispersed across studies with different objectives and without direct within-sample comparison (Mejbel et al., 2021; Thomson-Laing et al., 2025).

Fish eDNA is often present at very low concentrations in marine sediments (Huston et al., 2023; Matisoo-Smith et al., 2008; Stager et al., 2015; Tsuji et al., 2022). These records thus provide an appropriate sample type to evaluate the relationship between metabarcoding PCR replicate detection proportion and ddPCR-based quantification of low-abundance target DNA (Capo et al., 2022; Mejbel et al., 2021). Marine sediments typically contain highly degraded DNA, low target copy numbers, and a background dominated by non-target taxa, making them particularly appropriate for testing the sensitivity and quantitative performance of molecular approaches.

Here, we compared species-specific ddPCR quantification with metabarcoding PCR replicate detection frequency and relative read abundance for Atlantic cod (*Gadus morhua*) and Atlantic herring (*Clupea harengus*) in sedaDNA samples extracted from Icelandic marine sediment cores spanning more than three millennia. We assessed how these metabarcoding metrics correlate with independent estimates of target DNA concentration and whether ddPCR can detect low abundance signals that metabarcoding may fail to capture. In addition, we evaluated whether temporal trends in fish sedaDNA abundance are consistent with independently reconstructed SST variability, assessing how environmental change might influence molecular signals.

## 2 Materials and methods

### 2.1 Study site and sedimentary ancient DNA extracts

Two sediment cores from the North Icelandic shelf were analysed, core PC019 and core GC01 (Figure 1). Core PC019 was collected with a piston core at 471m depth during the 2022 DY150 cruise (core identifier: DY150-NIS-A-PC019; coordinates: 66.551721°N, –17.700281°W) (Scourse et al., 2024). Core GC01 was collected with a gravity core at 301m depth during the 2006 Millennium B05-2006 cruise (sample ID: B05-2006-GC01; coordinates: 66.5015°N, – 19.50567°W) (Eiríksson et al., 2006). Both sediment cores were chronologically constrained through a combination of tephrochronology and radiocarbon dating, with full age-depth models presented in Holman et al., (2025b). Sediment subsampling and sedaDNA extraction were carried out as part of the study described in Holman et al. (2025a) and are summarised here. PC019 was subsampled every 2 cm between 23 and 275 cm core depth, with additional 1 cm sampling around two putative tephra layers, whereas GC01 was subsampled every 4 cm from the core top to 180 cm. Together, the records span approximately three millennia, from 1315 BCE to 1785 CE.

**Figure 1:**
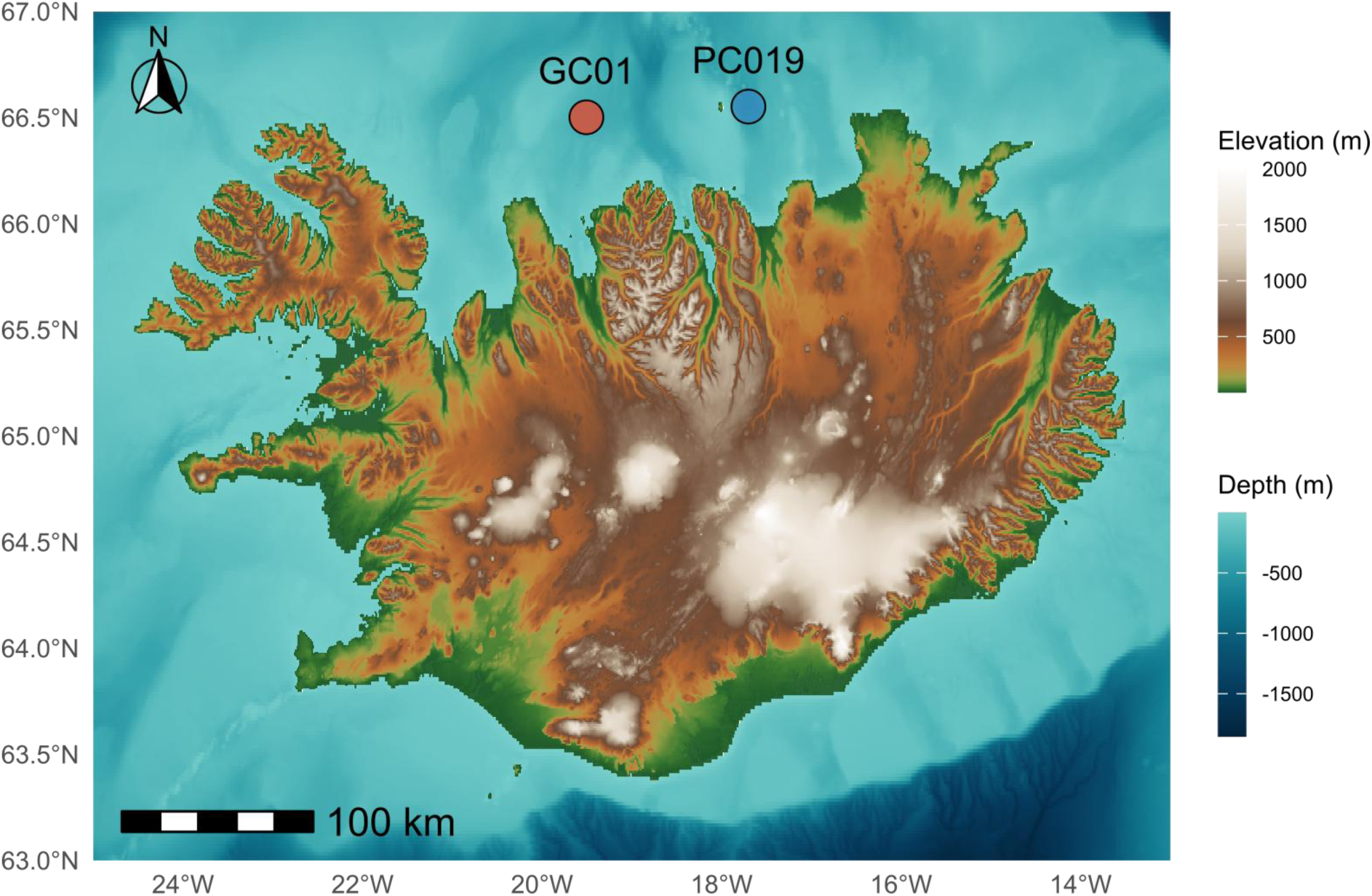
Map of Iceland showing the locations of the two marine sediment cores analysed in this study: GC01 (red) and PC019 (blue) on the North Icelandic shelf.

Environmental DNA extraction was conducted in dedicated ancient DNA clean laboratory facilities, following established ancient DNA protocols designed to minimise contamination risk (Heintzman et al., 2023; Knapp et al., 2012). A total of 158 sediment samples were extracted from PC019 and 46 samples from GC01 using the Qiagen MagAttract PowerSoil Pro DNA Kit (Version: October 2022). Extraction of 0.4-0.5 g of sediment per sample was carried out with minor changes to the manufacturer’s guidelines as outlined in Holman, et al. (2025a), which were specifically implemented to improve the recovery of short, degraded DNA fragments typical of sedaDNA. Extractions were processed in batches of 30, each including 29 sediment samples and one negative extraction control, resulting in a total of eight extraction blanks across the full dataset. DNA was eluted in 80 µL of solution C6 and stored at -20 °C until further analysis. For each sediment layer, a single DNA extraction was performed and subsequently used for both metabarcoding and ddPCR analyses.

### 2.2 Metabarcoding data

Metabarcoding PCR amplification and library preparation were carried out as part of the aforementioned study (Holman et al., 2025a). Here, we briefly summarise the relevant methodological details to provide context for the comparative analyses presented in this study.

All metabarcoding PCR preparation steps were conducted in dedicated ancient DNA facilities using pre-PCR clean laboratory protocols. A ca.130 bp fragment (excluding primers) of the nuclear 18S ribosomal RNA gene was targeted using a primer set designed to broadly amplify eukaryotic taxa, consisting of forward primer Euk1391f (Amaral-Zettler et al., 2009) and reverse primer EukBr (Stoeck et al., 2010) (Table S1). The decision to target the 18S RNA gene region was based on the parallel study (Holman et al., 2025a) that aimed to evaluate overall eukaryotic community composition. Both forward and reverse primers were labelled at the 5’ end with unique 8-mer nucleotide tags (>3 nucleotide differences between tags), each tag combination was used only once per amplicon pool to minimise tag-jumping artefacts. Eight PCR replicates were generated for each of the 204 eDNA extracts and at least two negative PCR controls were included every 40 PCR amplifications of sample DNA extracts. The 24 amplicon pools were built into sequencing libraries using the Tagsteady protocol as outlined in Carøe & Bohmann, (2020). One negative library control was incorporated into library preparation and sequenced alongside experimental libraries. The amplicon pool libraries were sequenced (250 bp paired-end) across two SP lanes on an Illumina NovaSeq 6000 instrument with a 10% PhiX spike-in.

Post-sequencing processing followed Holman et al. (2025a). Briefly, PCR replicates were demultiplexed using Cutadapt (v4.2) (Martin, 2011) and processed using DADA2 (v1.29.0) (Callahan et al., 2016) in R (v.4.2.2) (*R Core Team, 2025*) using default settings, except that reads were filtered with filterAndTrim (maxN = 0, maxEE = c(1,1), truncQ = 2). Amplicon sequence variants (ASVs) were inferred, chimeras removed, and replicate ASV tables merged. ASVs detected in only a single PCR replicate across the complete dataset (all samples and PCR replicates combined) were excluded as a standard metabarcoding quality-control step to reduce the influence of low-frequency artefacts, including sequencing errors, tag-jumping and stochastic amplification events (Agersnap et al. 2022; Jensen et al., 2022). Sequences outside the expected 70–170 bp range (excluding primers) were discarded. Data from both sequencing lanes were merged by collapsing ASVs with 100% identity and summing their read counts.

The ASVs were taxonomically assigned using *blastn* (v.2.12.0+) (Camacho et al., 2009) against the NCBI *nt* database (downloaded 27 Jan 2022) to return 200 hits (*-num_alignments* 200) per ASV; these were then parsed using a custom R script that uses a lowest common ancestor approach to assign a taxonomy to each ASV (*ParseTaxonomy*, doi:10.5281/zenodo.4671710). ASVs assigned to Actinopterygii (ray-finned fishes) were manually verified against the online NCBI blast portal (last accessed Jan 2024) following the taxonomic assignments to genus (Holman et al., 2025a). For comparison with ddPCR results, ASV counts were summed per PCR replicate across the genera *Gadus* and *Clupea* to represent read counts for cod and herring, respectively. Aggregation at the genus level was used to improve comparability with species-specific ddPCR results while avoiding overinterpretation of taxonomic assignments from the short 18S fragment. Although ddPCR assays targeted species-specific mitochondrial markers, the broader 18S metabarcoding marker did not consistently provide sufficient taxonomic resolution for confident species-level assignment across all fish taxa. Aggregating detections at the genus level therefore provided a more conservative framework for comparison between approaches. Although *Gadus* includes three species globally and *Clupea* includes two recognised species, Atlantic herring and Atlantic cod are the dominant representatives in the North Icelandic shelf region.

Metabarcoding data were summarised in two complementary ways: (1) Relative read abundance per genus was calculated separately for each PCR replicate as the proportion of reads assigned to target ASVs relative to the total number of reads in that replicate. Relative read abundance values were then averaged across the eight technical replicates to obtain a single per-sample estimate. (2) Detection frequency was expressed as the proportion of positive PCR (out of eight) in which the target taxon was detected. Per-sample relative read abundance values were used in downstream analyses, and overall means reported in the results represent the arithmetic average across all 204 sediment samples from both cores.

### 2.3 ddPCR assay design and implementation

Two species-specific ddPCR assays were used to target mitochondrial Cytochrome b DNA fragments for Atlantic cod (*Gadus morhua*) and Atlantic herring (*Clupea harengus*). The assays amplify fragments of 81 bp for Atlantic cod and 88 bp for Atlantic herring, respectively. These assays were previously developed and validated in Knudsen et al., (2019) using conventional qPCR with a dilution series of diluted PCR products from tissue samples to determine primer and probe efficiency and specificity. Primer and probe sequences for each assay are provided in Table S1. However, while the cod assay was designed to be species-specific, cross-amplification with closely related species cannot be entirely excluded. In particular, the assay was not tested against Greenland cod (*Gadus ogac)* or Alaska pollock (*Gadus chalcogrammus).* This potential taxonomic ambiguity is therefore unlikely to affect the temporal patterns analysed here but is considered when interpreting absolute species identity.

From the 204 extracted sedaDNA samples, ddPCR analyses were performed on a subset of 136 sediment extracts (93 from PC019 and 43 from GC01) selected based on sufficient remaining DNA volume and to capture the range of metabarcoding results observed across samples. Selection was not based on specific time intervals, as our focus here is more on the comparison of the molecular methods, and samples spanned the full temporal extent of both sediment records. The apparent clustering of some PC019 samples reflects the higher and uneven sampling resolution of the original metabarcoding dataset described in Holman et al. (2025a). The ddPCR analyses were performed on the same DNA extracts previously used for metabarcoding analyses (Holman et al., 2025a). In total, 109 samples were analysed for Atlantic cod and 109 for Atlantic herring, with an overlap of 82 samples analysed for both species. The resulting dataset therefore comprised 136 unique sediment extracts across both assays.

In total, 36 no-template controls (NTCs) were included across three ddPCR plates (12 per plate). NTCs were distributed across different rows and columns to monitor cross-well contamination and droplet misassignment. For each target species, two independent serial dilution series (10⁻² to 10⁴) of quantified PCR products derived from tissue samples (Knudsen et al., 2019) were included to assess assay linearity and dynamic range. Although ddPCR provides absolute quantification based on Poisson statistics and does not rely on external standard curves, dilution series were used here to verify assay performance and consistency across plates.

All ddPCR reactions were prepared in a dedicated pre-PCR laboratory under strict contamination-control procedures. Reaction set-up was conducted in laminar flow hoods, with surfaces and consumables routinely decontaminated using bleach, ethanol and UV irradiation. Droplets were generated using the Bio-Rad QX200™ Droplet Generator. Thermal cycling conditions were: 95 °C for 10 min; 40 cycles of 94 °C for 30 s and 60 °C for 1 min; followed by 98 °C for 10 min. Droplet fluorescence was measured using the Bio-Rad QX200™ Droplet Reader. Reaction compositions and final concentrations are provided in Table S2. Each sediment extract was analysed once per target assay due to limited extract volume. Large reaction volumes were used to maximise target molecule recovery from low-concentration sedaDNA extracts.

Raw fluorescence amplitude data were processed in Bio-Rad QX Manager Software (v2.3). Thresholds separating positive and negative droplets were manually adjusted for each assay based on amplitude histograms, placing the threshold at the trough between clearly separated droplet populations (Witte et al., 2016) for each plate separately. This approach ensured consistent discrimination between true positive droplets and background noise. Fluorescence amplitudes were inspected across dilution series to identify and exclude ambiguous “star” droplets (Porco et al., 2022), allowing species-specific thresholds to be defined.

Absolute concentrations (copies µL⁻¹) were calculated directly from positive and total droplet counts using Poisson-based concentration estimates implemented in Bio-Rad QX Manager Software (v2.3). Linear regression models used to assess assay efficiency and dynamic range were fitted exclusively to the four highest-concentration dilutions, corresponding to values above the inferred limit of quantification (LOQ) established for the original qPCR assays (Knudsen et al., 2019). For comparative analyses with metabarcoding data, ddPCR results were expressed as absolute copy numbers per µL of PCR performed on extracted sedaDNA.

### 2.4 Climatic data

Previously published climatic records were sourced from a sediment core (MD99-2275, 66.551667 -17.699833) collected in 1999 within 100 m of the sampling site of PC019, a Holocene record covering approximately the last 9300 years (Jiang et al., 2015), to represent local sea surface temperature (SST) variability on the North Icelandic shelf. These records consisted of two SST proxies, a diatom transfer function (Jiang et al., 2015) and C37 alkenones (Sicre et al., 2021). For each record 100-year splines were generated using the *detrend.series* function from the *dplR* package (v1.7.6.) (Bunn, 2010) in R v4.3 (*R Core Team, 2025*). Climatic values for each sediment layer were extracted from these splines.

### 2.5 Data analysis and statistical approaches

To explore non-linear temporal patterns of ddPCR concentration within individual sediment cores, generalised additive models (GAMs) were fitted using the *gam* function from the mgcv package (v1.9-0). These models evaluated the relationship between ddPCR concentration and sample age and were specified as: [ddPCR (copies µL^-1^)] = s(Age, k = 10)+ε. Where s() denotes a spline smoothing function with a maximum of 10 basis functions (k = 10) fitted using Restricted Maximum Likelihood (REML).

Following detection of positive droplets in NTC wells, the mean NTC concentration for each assay was calculated. Experimental samples with concentrations below this threshold were excluded from downstream analyses.

To examine associations between ddPCR-derived DNA concentration and complementary molecular and environmental variables, we fitted ordinary least-squares linear models using the *lm* function from the stats package (v3.6.2). Models were run separately for each target species and also with target species included as an additive predictor. Predictor variables included (1) metabarcoding PCR replicate detection frequency, (2) metabarcoding relative read abundance (based on ASV sequence counts), (3) reconstructed sea surface temperature (SST), and (4) sediment core location.

All statistical analyses were performed in R (v4.3.1) and statistical significance was evaluated at p ≤ 0.05.

## 3 Results

### 3.1 Droplet digital PCR

Fluorescence amplitude distributions across the standard dilution series (Figure S3), supported an upper fluorescence threshold of 7,000 amplitude units for Atlantic cod and 13,000 amplitude units for Atlantic herring to distinguish positive from negative droplets. Dilution series showed strong linearity (R² > 0.99) for both the Atlantic cod and Atlantic herring assays across concentrations above the previously established qPCR limit of quantification (LOQ) (Knudsen et al., 2019; Figure S2).

Across the 136 analysed samples, ddPCR detected Atlantic cod in 109 samples and Atlantic herring in 110 samples. Both Atlantic cod and Atlantic herring were jointly detected in 82 of these samples. Among samples with positive detections, concentrations ranged from 0.12 to 21.37 copies µL⁻¹ for Atlantic cod and from 0.42 to 45.31 copies µL⁻¹ for Atlantic herring (Figure 2). No-template controls (NTCs) showed low-level amplification across the workflow for both target species, with 23 of 24 Atlantic cod controls and all 24 Atlantic herring controls showing positive detections, although signal intensities remained low (Atlantic cod: 0-8 copies µL^-1^; Atlantic herring: 0.18 - 4.30 copies µL^-1^) (Figure S4). These values represent the original concentrations observed across NTCs and were used to define species-specific filtering thresholds. Samples with concentrations below the mean concentration observed among NTCs for each species were excluded from subsequent analyses.

**Figure 2:**
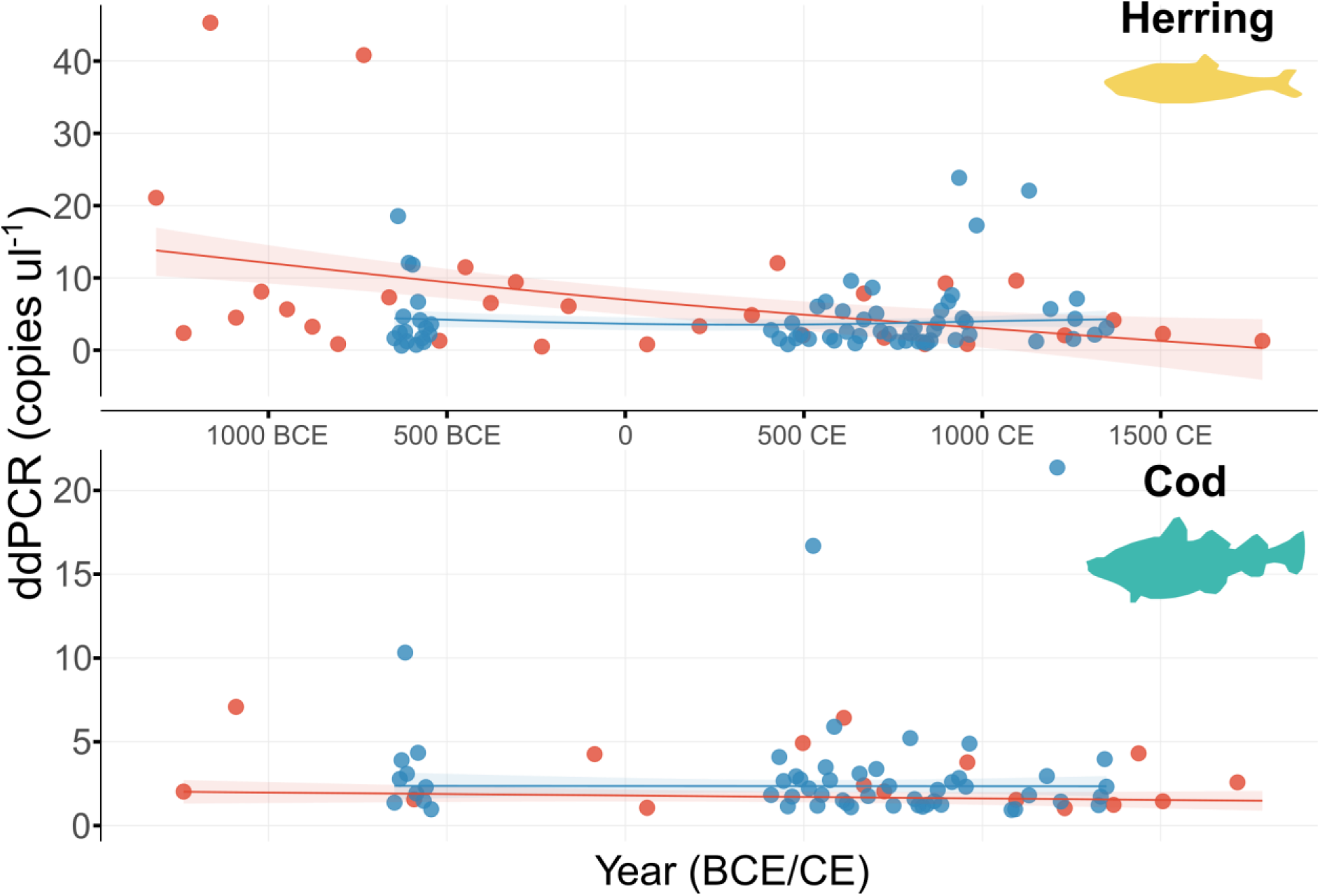
ddPCR copy numbers per microliter plotted over time for each core and species. Generalised additive models (GAMs) predictions are shown as solid lines, with shaded areas representing 95% confidence intervals.

Generalised additive models (GAMs) fitted independently for each core and species showed no clear pattern in DNA concentration through time except in GC01 for Atlantic herring (Figure 2). In this case, the GAM indicated a significant relationship between ddPCR concentration and age (GAM; F = 3.36, p = 0.047; Table S3), with the fitted smooth suggesting a decline in herring sedaDNA concentration across the record.

### 3.2 Metabarcoding data and comparison with ddPCR quantification

A total of six amplicon sequence variants (ASVs) from metabarcoding were taxonomically assigned to *Clupea*, with an average relative read abundance of 1.35x10^-4^ (± s.d. 2.87x10^-4^) per PCR replicate. *Clupea* was detected in 192 of 204 sediment extracts (94.1%). Two ASVs were assigned to *Gadus*, yielding an average of around 8.32x10^-5^ (± s.d. 1.89x10^-4^) relative read abundance for each PCR replicate. *Gadus* was detected in 193 of 204 sediment extracts (94.6%). Beyond the target genera, fish-associated ASVs included representatives from four additional families (Ammodytidae, Myctophidae, Pleuronectidae and Salmonidae), one additional genus (*Melanostigma*), and one order-level assignment (Perciformes). Many detections could only be resolved to higher taxonomic levels, reflecting the limited taxonomic resolution of the short 18S marker. Overall, fish reads represented 0.05% of the total metabarcoding dataset (126,528 fish reads from 244,934,846 total reads), suggesting that fish DNA constituted only a small component of the overall eDNA pool.

Models including all samples showed a significant positive relationship between log_10_-transformed ddPCR-quantified DNA copy number and metabarcoding PCR detection frequency at the genus level (F = 16.3, p < 0.001; Figure S4, Table S4). In species-specific models, this relationship was significant for Atlantic herring (F = 4.73, p = 0.032) but not for Atlantic cod (F = 1.05, p = 0.308; Figure S4, Table S4). The analysis was then repeated after applying species-specific NTC thresholds to evaluate whether the observed relationships remained robust following exclusion of low-concentration signals close to background levels (Figure 3). After application of species-specific NTC thresholds, samples with ddPCR concentrations below the threshold for a given target species were excluded from the corresponding species-specific analysis. Following filtering, 109 unique sediment extracts were retained across both assays (74 from PC019 and 35 from GC01), including 64 samples retained for Atlantic cod and 92 for Atlantic herring, with an overlap of 46 samples retained for both species. The positive relationship remained significant at the genus level (F = 8.976, p < 0.001; Figure 3, Table S4) and was significant in both species-specific models for Atlantic cod (F = 6.78, p = 0.012) and Atlantic herring (F = 4.58, p = 0.035).

**Figure 3:**
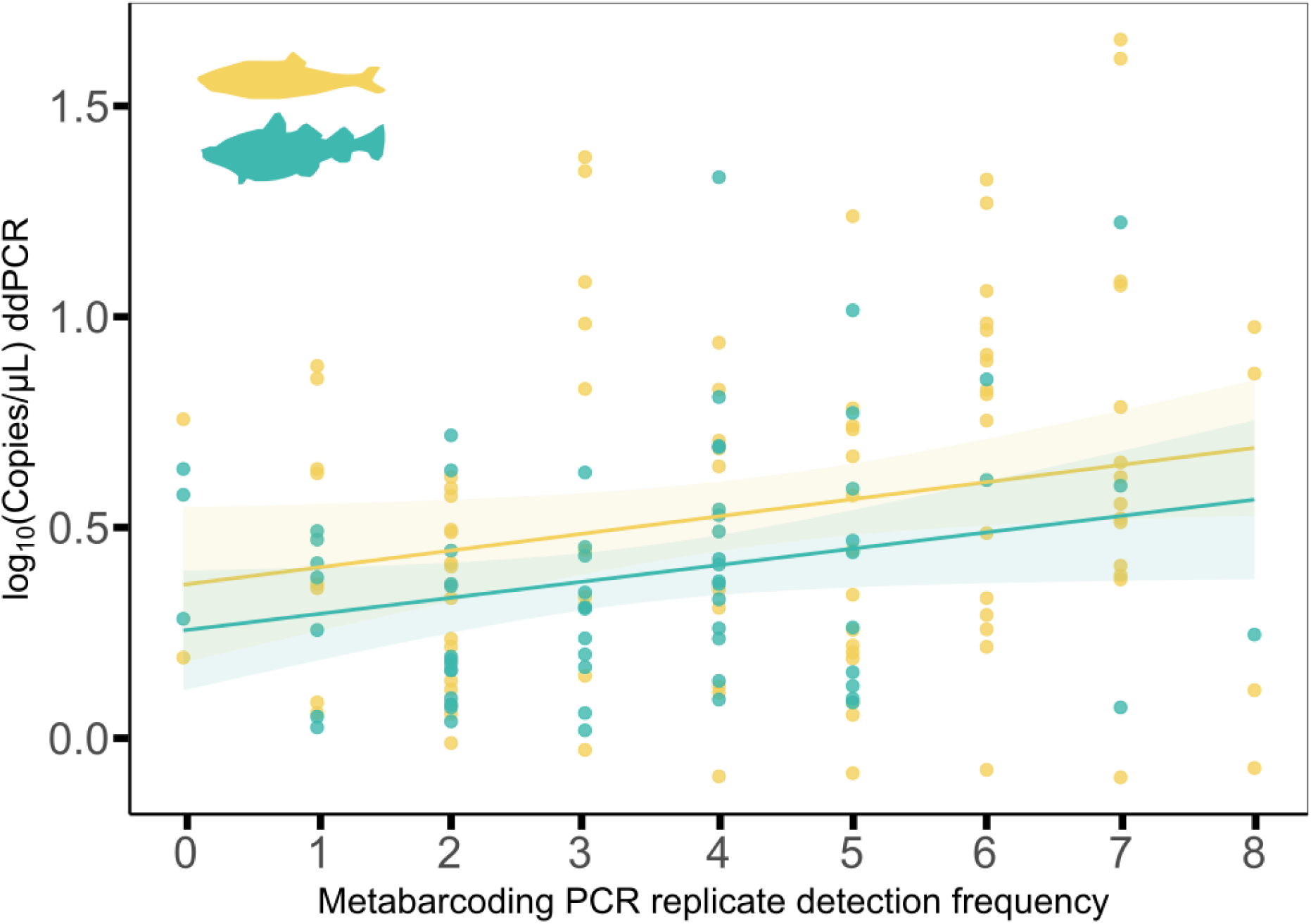
Relationship between ddPCR concentration and metabarcoding PCR replicate detection frequency for genus Clupea (yellow) and Gadus (turquoise) across sediment samples from two Icelandic marine sediment cores (PC019 and GC01; data combined). All ddPCR concentrations (copies µL⁻¹) are shown on a log_10_ scale and plotted against the number of positive metabarcoding PCR replicates per sediment sample (out of eight PCR replicates per sample). Solid lines show linear regression fits with shaded 95% confidence intervals.

When metabarcoding relative read abundance was used as predictor, ddPCR concentrations were positively associated with read abundance for Atlantic cod (F = 5.63, p = 0.021; Figure 4, Table S4) and for Atlantic herring (F = 5.97, p = 0.016). Finally, metabarcoding relative read abundance was strongly correlated with PCR replicate detection frequency for both genera, *Gadus* (F = 41.29, p < 0.001) and *Clupea* (F = 266.60, p < 0.001; Figure S5, Table S4).

**Figure 4:**
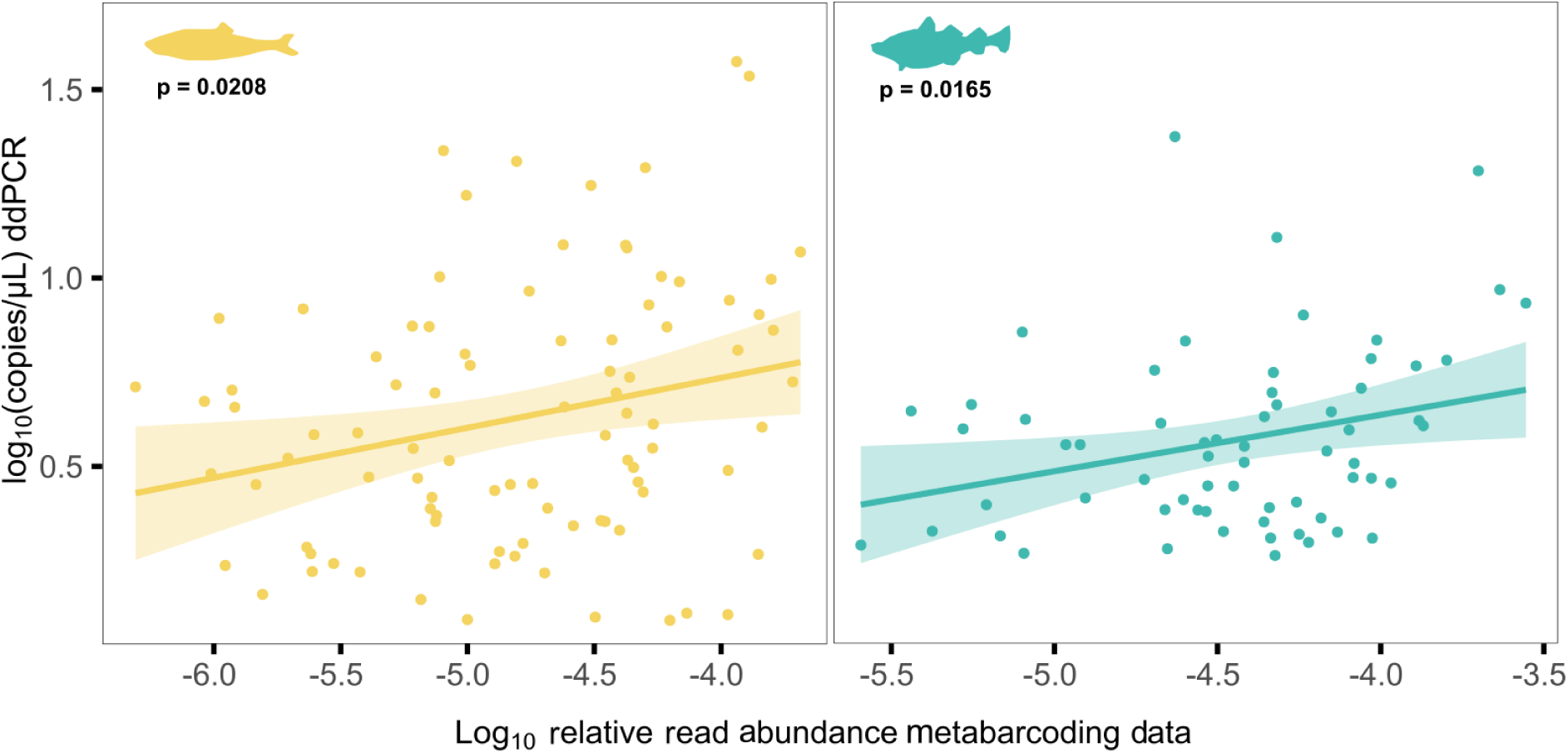
ddPCR concentration versus metabarcoding relative read abundance on a log10-log10 scale for Clupea harengus (yellow) and Gadus morhua (turquoise). Statistical significance (p-values) of the correlations is indicated on each plot.

### 3.3 Relationship between ddPCR quantification and environmental conditions

To explore potential environmental drivers of fish sedaDNA variability, ddPCR-derived DNA concentrations were compared with reconstructed sea surface temperature (SST) records from the North Icelandic shelf (Figure 5). Atlantic herring showed a positive association with reconstructed environmental conditions, for alkenone-based SST estimates (F = 5.4, p = 0.022) but a non-significant trend for SST estimates derived from a diatom transfer function (F = 3.31, p = 0.072) (Figure 5; Table S4). No significant relationships between ddPCR concentration and SST were observed for Atlantic cod (Table S4).

**Figure 5:**
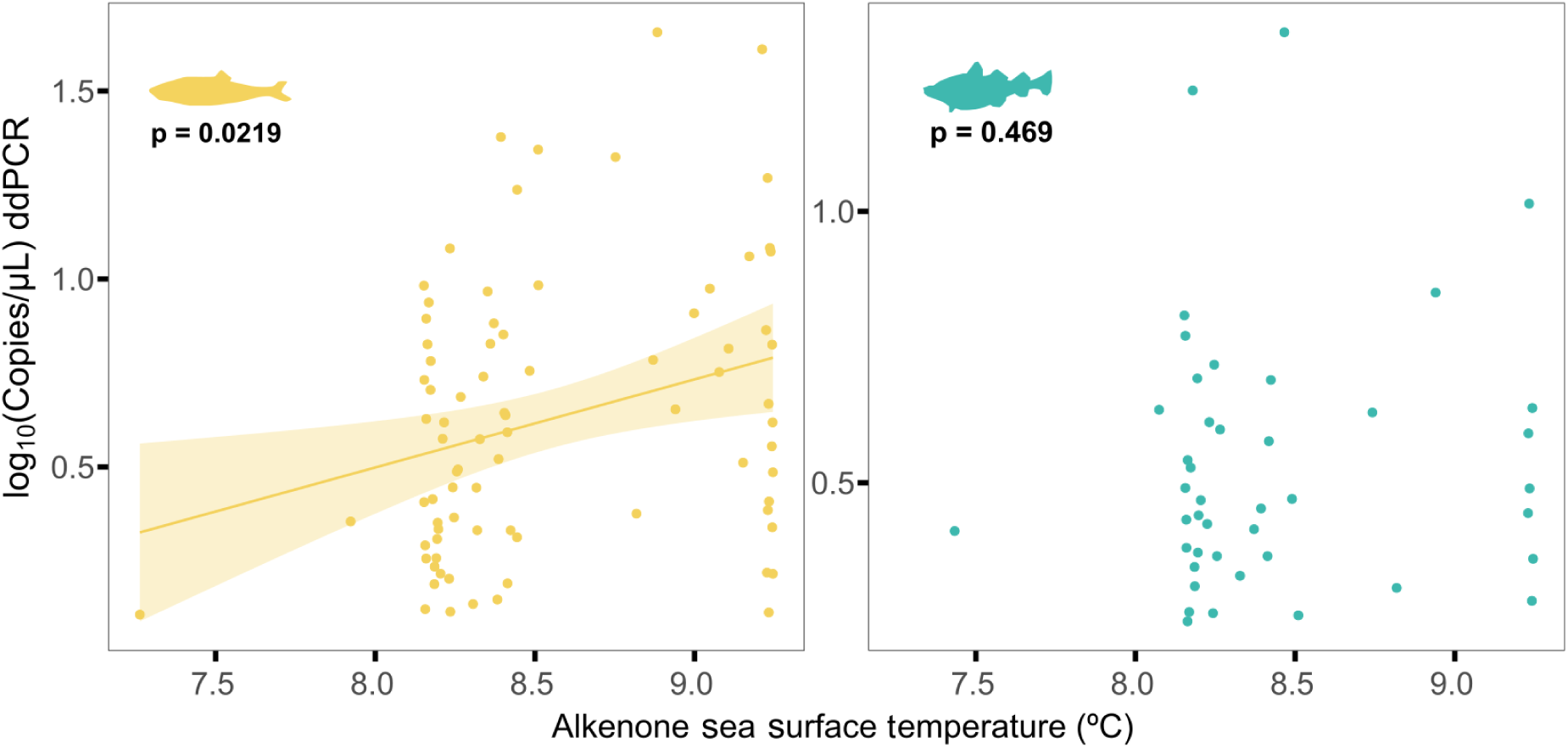
Log10-transformed ddPCR sedaDNA concentrations plotted against reconstructed alkenone-derived sea surface temperature (SST) estimates for Atlantic herring (Clupea harengus, yellow) and Atlantic cod (Gadus morhua, turquoise).

## 4 Discussion

Different methods capture distinct aspects of biodiversity from eDNA samples. Through directly comparing ddPCR and metabarcoding of fish sedaDNA from Icelandic marine sediment records, we show that when eDNA is rare, both approaches contain information about target DNA abundance. Although metabarcoding data typically provides a compositional description of a community, our results suggest that more might be gleaned from these datasets. Our comparison of molecular methods arose from a project that assessed the eukaryotic community composition in the core samples using an 18S gene region (Holman et al., 2025a). While this region proved imperfect for vertebrate species here we show that even limited data carries valuable information about past ecosystems.

### 4.1 Linking metabarcoding PCR replicate detection frequency to absolute ddPCR quantification

A growing number of studies report the proportion of positive metabarcoding PCR replicates for a given taxa (Clarke et al., 2024; Garcés-Pastor et al., 2022; Holman et al., 2025b; Zetter et al., 2025), implicitly suggesting that this metric reflects underlying target DNA concentration or relative signal strength. However, evidence supporting this interpretation has largely remained indirect, often relying on correlations with independent ecological or environmental proxies (Alsos et al., 2020; Holman et al., 2025a; Zetter et al., 2025). Here, we show a significant positive correlation between ddPCR-derived DNA concentration and the proportion of positive metabarcoding PCR replicates per sediment sample (Figure 3 & Figure S4), providing empirical support to interpret PCR replicate detection frequency as a semi-quantitative proxy for relative target DNA concentration in metabarcoding-based sedaDNA studies.

The utility of PCR replicate detection frequency as a quantitative proxy is likely constrained to contexts where target DNA is present at low concentrations and detection is probabilistic (Chen & Ficetola, 2020). In the present study, most ddPCR concentrations were below approximately 20 copies µL^-1^, within a range where replicate detections varied substantially among samples rather than all replicates being positive or negative. At higher DNA concentrations, detection across PCR replicates is expected to saturate (i.e. approach 100% positive replicates) (Bista et al., 2017; Clarke et al., 2019; Elbrecht et al., 2017; Salter et al., 2019). In such cases, PCR replicate detection frequency may provide little additional quantitative information beyond presence–absence, and relative read abundance or absolute quantification approaches may be more informative (Doi et al., 2015; Elbrecht & Leese, 2015).

Overall, these results indicate that PCR replicate detection frequency is most informative for low-abundance sedaDNA targets, where stochastic amplification dominates detection outcomes and replicate-based metrics retain discriminatory power (Alsos et al., 2020; Chen & Ficetola, 2020). Consequently, replicate detection frequency should be interpreted as a semi-quantitative metric rather than a universally linear indicator of target DNA concentration. In this context, ddPCR and metabarcoding provide complementary information from the same sedaDNA extracts: ddPCR offers sensitive, target-specific quantification at both low and high copy numbers (Guri et al., 2024; Pinheiro et al., 2012), while replicated metabarcoding detections reflect probabilistic detection patterns associated with variation in target DNA concentration (Clarke et al., 2019).

### 4.2 Detecting fish eDNA using ddPCR and metabarcoding

Fish eDNA can remain detectable in marine and freshwater sediments for thousands of years (Kuwae et al., 2020; Lopez et al., 2024). Previous studies have used metabarcoding PCR replicate detection frequency to track fish sedaDNA signals without explicit calibration against independent abundance estimates (Huston et al., 2023; Rossouw et al., 2025). Here we showed that both ddPCR and 18S metabarcoding data detected Atlantic cod and Atlantic herring across more than three millennia. Although the correlation between methods was low (R^2^= 0.105; Figure 3, Table S4), both fish species were detected consistently across both cores. ddPCR concentrations showed structured temporal variation through time (Figure 2) that partially overlapped with metabarcoding patterns previously reported in Holman et al. (2025a). An important difference between ddPCR and metabarcoding in this study is taxonomic resolution: ddPCR targeted species-specific mitochondrial fragments, whereas metabarcoding detections were aggregated at the genus level because the short 18S marker did not consistently support confident species-level assignment. Detections in the analysed marine sediment samples could, in principle, also originate from congeners, for example *Gadus macrocephalus*, which has been reported from northern Greenland (Møller et al., 2010) and may have had a more southern distribution in the past. However, given the strong ecological and biogeographic dominance of Atlantic cod on the North Icelandic shelf throughout the study period (Eiríksson et al., 2006; Ólafsdóttir et al., 2021), and the absence of evidence for persistent local populations of other *Gadus* species in this region (Rose, 2007), detections are most parsimoniously interpreted as Atlantic cod. Clear differences between species were observed through time, most notably the decline in Atlantic herring sedaDNA concentration in core GC01. The positive relationship between herring ddPCR concentration (Figure 3) and both metabarcoding relative read abundance (Figure S5) and reconstructed sea surface temperature (SST) (Figure 5), add support for sedaDNA-based tracking of environmentally structured variation in species occurrence. In contrast, no comparable relationships with SST were detected for Atlantic cod, suggesting differences in how sedaDNA signals from the two species integrate ecological variability through time. These differences may reflect contrasting sedaDNA deposition and preservation (Corinaldesi et al., 2008) as a result of the different eDNA profiles (Brandão-Dias et al., 2025b) of Atlantic cod (demersal) and Atlantic herring (pelagic). Life history, behaviour, and habitat use all likely influence where and how DNA enters the sedimentary environment, influencing signal strength and environmental sensitivity (Harrison et al., 2019; Port et al., 2016).

Assuming the observed differences in sedaDNA concentration, at least in part, reflect underlying species abundance, the significant relationship between SST and Atlantic herring sedaDNA concentrations suggests climate-related shifts in herring occurrence on the North Icelandic shelf (Figure 5), consistent with previous evidence of temperature sensitivity in this species (Campana et al., 2020; Holman et al., 2025a; Rose, 2005; Trochta et al., 2020). In contrast, Atlantic cod showed no significant relationship between SST and ddPCR-derived sedaDNA concentrations, possibly reflecting its broader ecological tolerance and more complex trophic and life-history dynamics (Astthorsson et al., 2007; Giebner et al., 2020; Martínez-García et al., 2021; Ólafsdóttir et al., 2021). However, alternative explanations should also be considered, including differences in how species-specific sedaDNA signals integrate ecological variability over time or potential taxonomic resolution limits in distinguishing *Gadus* species within the sedimentary record for each approach. The SST reconstructions are derived from a nearby (ca. 100 meters away) sediment core, and thus may not fully capture broader variation in environmental conditions driving fish eDNA abundance. While SST is only one of many environmental variables structuring fish distributions in the absence of alternatives it is a valid parameter to contextualise if molecule counts obtained through metabarcoding and ddPCR are measuring a real ecological response in the past. .

### 4.3 Methodological constraints and sources of stochasticity in low-abundance sedaDNA

Here, PCR stochasticity refers to random variation in detection and quantification from probabilistic sampling of target molecules. In ddPCR, this primarily reflects random partitioning of DNA molecules into droplets, leading to variable detection at very low template concentrations (Pinheiro et al., 2012). Detection sensitivity further depends on assay design, including amplicon length, probe–target affinity, and amplification efficiency. In our study, ddPCR assays targeted short mitochondrial fragments (81–88 bp), whereas the 18S metabarcoding fragment was approximately 130 bp. Shorter amplicons are generally more likely to amplify successfully from highly degraded sedaDNA templates, which may partly explain differences in detection consistency between ddPCR and metabarcoding results (Brandão-Dias et al., 2025a; West & Deagle, 2025).

Metabarcoding, in contrast, is affected by additional sources of stochasticity, including variability among PCR replicates, PCR amplification of many different targets within the same reaction, and constraints imposed by sequencing depth (Elbrecht & Leese, 2015; Nichols et al., 2018). Compositional effects within the total DNA pool may further influence amplification dynamics, as highly abundant taxa can dominate sequencing output and reduce detectability of rare target sequences. Although emerging PCR-free approaches may reduce some amplification-related biases (Urban et al., 2021), their applicability to highly degraded, low-template sedaDNA has yet to be demonstrated. Despite these limitations, metabarcoding provides broader ecological context through the detection of other taxa and can retain informative presence–absence signals even information when ancient DNA signals approach analytical detection limits (Alsos et al., 2020; Clarke et al., 2019; Holman et al., 2025a).

The relatively weak relationships observed between methods suggest that additional factors, including differences in marker type (mitochondrial versus nuclear DNA), taxonomic resolution and differences in molecule partitioning and technical replication between approaches may also contribute to the observed scatter (Deiner et al., 2017; Elbrecht & Leese, 2015; Nichols et al., 2018). Although the observed relationships explained only a modest proportion of variance, the consistent positive associations across analyses nevertheless indicate that metabarcoding replicate detection frequency retains quantitative information related to underlying target DNA availability.

The observed changes in statistical significance following exclusion of samples with ddPCR concentrations below the NTC-derived threshold indicate that relationships between methods may be sensitive to filtering decisions when target signals occur close to detection limits. This is typical of low-abundance sedaDNA datasets, where the ecological signal is often close to analytical limits (Chen & Ficetola, 2020; Clarke et al., 2019). However, the significant genus-level relationship remained after filtering (Figure 3), indicating that the overall pattern was consistent across analytical approaches.

The observed correlation between ddPCR concentrations and metabarcoding detection frequency likely reflects the shared stochastic sampling process during PCR. Under low-template conditions, small differences in the number of target molecules entering individual droplets or PCR reactions can result in variable detection outcomes among samples, even when underlying DNA concentrations are similar. Stochasticity is therefore inherent to low abundance sedaDNA analyses and, when explicitly considered, may improve quantitative interpretation rather than simply represent methodological noise.

## Acknowledgments

This research was supported by the European Research Council (ERC) through the SeaChange project (Grant agreement No. 856488). Elena Baños Lara was additionally supported by the iMOVE (Grant agreement No. IMOVE24269) international mobility programme of the Spanish National Research Council (CSIC) during her research stay at the University of Copenhagen, the Spanish Ministry of Science, Innovation and Universities grants PID2023-146307OB-C22 (MICIU/AEI/10.13039/501100011033 and FEDER, UE) and PID2020-118550RB (MICIU/AEI/https://doi.org/10.13039/501100011033), and the grant TED2021-132228B-C22 funded by MCIN/AEI/10.13039/501100011033 and the European Union (“NextGenerationEU”/PRTR). Steen Wilhelm Knudsen was supported through the OBAMA-NEXT (Observing and Mapping Marine Ecosystems – Next Generation Tools) project, funded by the European Union under the Horizon Europe program (grant agreement no. 101081642), www.obama-next.eu. We are grateful to the entire SeaChange team for providing access to sedaDNA extracts and for constructive discussions throughout the study.

## Author contributions

Elena Baños and Luke E. Holman conceived the study, designed the methodology, conducted laboratory work, and performed data analysis and interpretation. Steen W. Knudsen contributed to the experimental design and optimization of ddPCR protocols. All authors contributed to writing and revising the manuscript and approved the final version for submission.

## Data availability statement

Data available via DOI https://doi.org/10.5281/zenodo.21284352 (Baños et al., 2026). The raw eDNA metabarcoding data can be found at the European Nucleotide Archive and processed data at PRJEB78865 and under DOI https://doi.org/10.5281/zenodo.15575942 (Holman et al., 2025c).

## Statement on inclusion

This study analysed previously generated DNA extracts from Icelandic marine sediment cores that were provided as archived material. No new field sampling, laboratory work, or research activities were conducted in Iceland for this study. The research involved no human participants, animals, or Indigenous knowledge, and therefore did not require ethical approval. Original sample collection and export were conducted under the permits and regulations applicable at the time of sampling. Relevant literature from Icelandic researchers was consulted in interpreting the results.

## Supporting Information

**Table S1:**
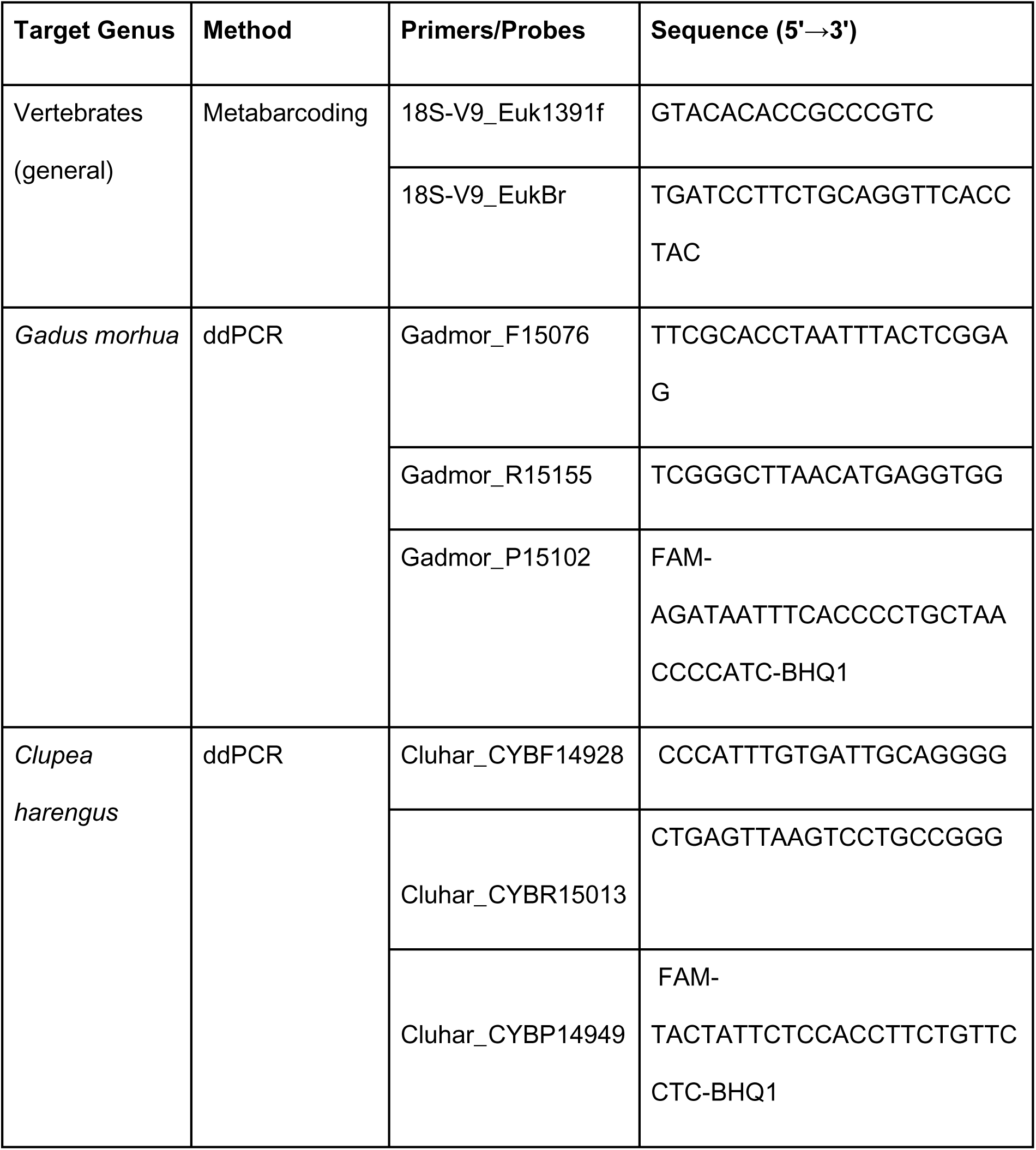
Primers and probes used in this study for both metabarcoding and ddPCR assays. Metabarcoding primers target the V9 region of the 18S rRNA gene, while species-specific ddPCR primers and probes were used to quantify mitochondrial DNA from *Gadus morhua* and *Clupea harengus*. All sequences are listed in 5’→3’ orientation.

**Table S2:** Reaction composition and final concentrations used for the *Clupea harengus* (herring) and *Gadus morhua* (cod) droplet digital PCR (ddPCR) assays. Each 22 µL reaction consisted of Bio-Rad ddPCR Supermix for Probes (no dUTP), species-specific forward and reverse primers, hydrolysis probe, molecular-grade water, and 8 µL of DNA template. Primer and probe sequences are listed in Knudsen et al. (2019). Final concentrations were optimised based on primer efficiency tests and correspond to the concentrations used for all sedaDNA extracts in this study. Reaction mixes were prepared in a dedicated clean laboratory using single-use consumables and negative controls (NTCs) included on each plate.

| Target Species | | Primer / Probe | Volume (μL) | Final concentration |
| --- | --- | --- | --- | --- |
| ***Gadus morhua*** | Bio-Rad ddPCR Supermix | - | 11 | 1x |
|  | Primer forward | Gadmor_F15076 | 0,5 | 400 nM |
|  | Primer reverse | Gadmor_R15155 | 0,5 | 800 nM |
|  | Probe | Gadmor_P15102 | 0,5 | 200 nM |
|  | ddH<sub>2</sub>O | - | 1.5 |  |
|  | DNA template | - | 8 |  |
|  | Total volumen | - | 22 |  |
| ***Clupea harengus*** | Bio-Rad ddPCR Supermix | - | 11 | 1x |
|  | Primer forward | Cluhar_CYBF14928 | 0,5 | 200 nM |
|  | Primer reverse | Cluhar_CYBR15013 | 0,5 | 1000 nM |
|  | Probe | Cluhar_CYBP14949 | 0,5 | 200 nM |
|  | ddH<sub>2</sub>O | - | 1,5 |  |
|  | DNA template | - | 8 |  |
|  | Total volumen | - | 22 |  |

**Table S3:**
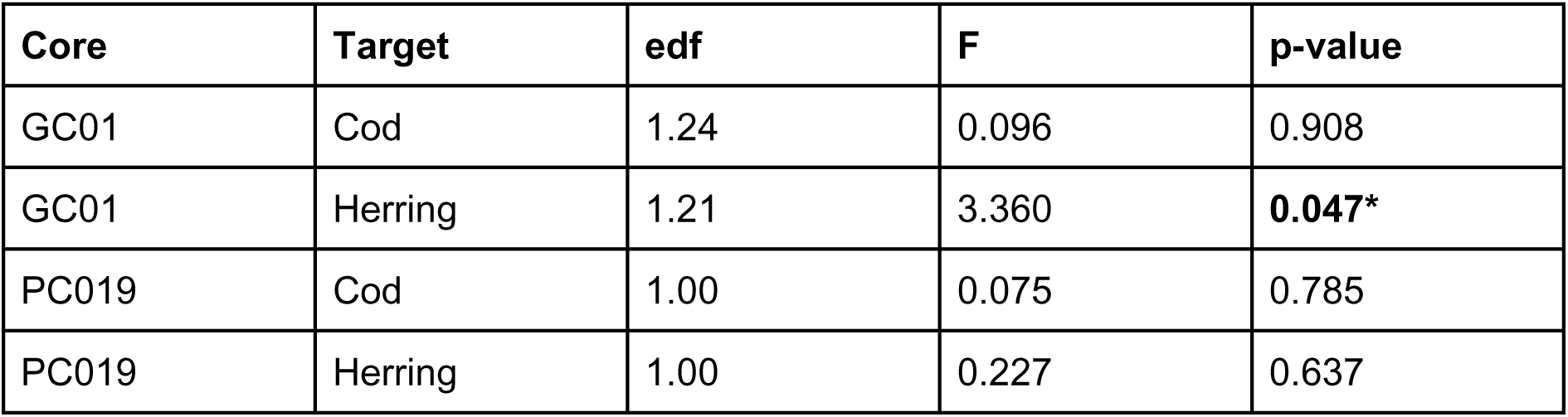
Summary of generalized additive models (GAMs) fitted to ddPCR concentrations as a function of time. Reported statistics correspond to the smooth term s(Date) for each Target × Core combination. edf indicates the estimated degrees of freedom of the smooth; values close to 1 indicate an approximately linear relationship. Significance was assessed using approximate F-tests. Significant effects (p < 0.05) are indicated by an asterisk (*).

| Core | Target | edf | F | p-value |
| --- | --- | --- | --- | --- |
| GC01 | Cod | 1.24 | 0.096 | 0.908 |
| GC01 | Herring | 1.21 | 3.360 | **0.047\*** |
| PC019 | Cod | 1.00 | 0.075 | 0.785 |
| PC019 | Herring | 1.00 | 0.227 | 0.637 |

**Table S4:** Summary of linear models testing the relationship between log_10_-transformed ddPCR concentrations (cleaned dataset) and metabarcoding detection metrics detection frequency (DetFreq; number of positive PCR replicates) and relative read abundance (RelAbund), as well as environmental variables, including sea surface temperature reconstructed from alkenone (SST-Alk) and diatom (SST-Dia) proxies. Results are shown separately for *Gadus morhua*, *Clupea harengus*, and combined models where applicable. Statistically significant p-values (p < 0.05) are indicated with an asterisk (*).

| Model | df | R<sup>2</sup> | R<sup>2</sup> adj | F | p-value | Std Er |
| --- | --- | --- | --- | --- | --- | --- |
| **COD:** *DetFreq ~ log<sub>10</sub>(ddPCR)*<br>(All samples) | 107 | 0.010 | <0.001 | 1.048 | 0.308 | 0.403 |
| **HERRING:** *DetFreq ~ log<sub>10</sub>(ddPCR)* (All samples) | 107 | 0.042 | 0.033 | 4.726 | **0.032\*** | 0.453 |
| **BOTH:**<br>*DetFreq ~ log<sub>10</sub>(ddPCR) + Target* (All samples) | 215 | 0.131 | 0.123 | 16.32 | **<0.001\*** | 0.428 |
| **COD:** *DetFreq ~ log<sub>10</sub>(ddPCR)*<br>(Clean samples) | 62 | 0.099 | 0.084 | 6.776 | **0.012\*** | 0.271 |
| **HERRING:** *DetFreq ~ log<sub>10</sub>(ddPCR)* (Clean samples) | 92 | 0.0467 | 0.038 | 4.584 | **0.035\*** | 0.398 |
| **BOTH:**<br>*DetFreq ~ log<sub>10</sub>(ddPCR) + Target* (Clean samples) | 153 | 0.105 | 0.093 | 8.976 | **<0.001\*** | 0.351 |
| **COD:** $\log_{10}(\text{RelAbund}) \sim \log_{10}(\text{ddPCR})$ (Clean samples) | 62 | 0.083 | 0.068 | 5.628 | **0.021\*** | 0.478 |
| **HERRING:** $\log_{10}(\text{RelAbund}) \sim \log_{10}(\text{ddPCR})$ (Clean samples) | 90 | 0.062 | 0.052 | 5.972 | **0.016\*** | 0.624 |
| **Gadus:** $\text{DetFreq} \sim \log_{10}(\text{RelAbund})$ | 62 | 0.400 | 0.390 | 41.29 | **<0.001\*** | 0.387 |
| **Clupea:** $\text{DetFreq} \sim \log_{10}(\text{RelAbund})$ | 90 | 0.748 | 0.745 | 266.6 | **<0.001\*** | 0.324 |
| **COD:** $\text{SST}(\text{Alk}) \sim \log_{10}(\text{ddPCR})$ (Clean samples) | 62 | 0.008 | -0.007 | 0.531 | 0.469 | 0.284 |
| **HERRING:** $\text{SST}(\text{Alk}) \sim \log_{10}(\text{ddPCR})$ (Clean samples) | 90 | 0.057 | 0.047 | 5.444 | **0.022\*** | 0.397 |
| **COD:** $\text{SST}(\text{Dia}) \sim \log_{10}(\text{ddPCR})$ (Clean samples) | 62 | 0.017 | 0.001 | 1.089 | 0.301 | 0.283 |
| **HERRING:** $\text{SST}(\text{Dia}) \sim \log_{10}(\text{ddPCR})$ (Clean samples) | 90 | 0.036 | 0.025 | 3.314 | **0.072** | 0.401 |

**Figure S1:**
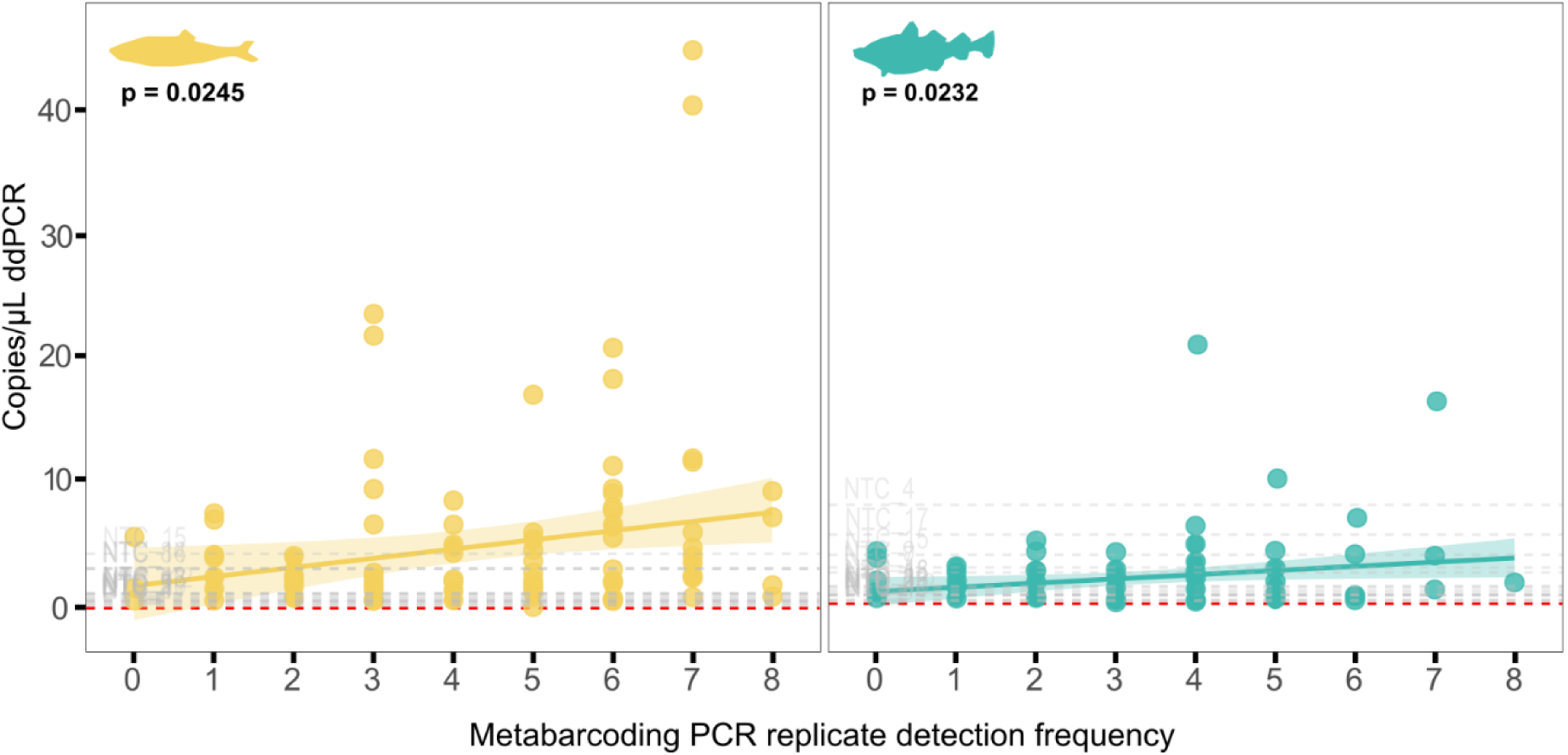
Total ddPCR concentrations against number of positive metabarcoding PCR replicates for Clupea harengus *(yellow) and* Gadus morhua *(turquoise). Negative control samples (NTCs) are shown in grey; red dashed lines indicate the species-specific mean concentration of NTCs. Linear regression model fits are shown by a continuous line with shaded 95% confidence intervals; statistical significance (p-values) for each fit is shown on each plot*.

**Figure S2:**
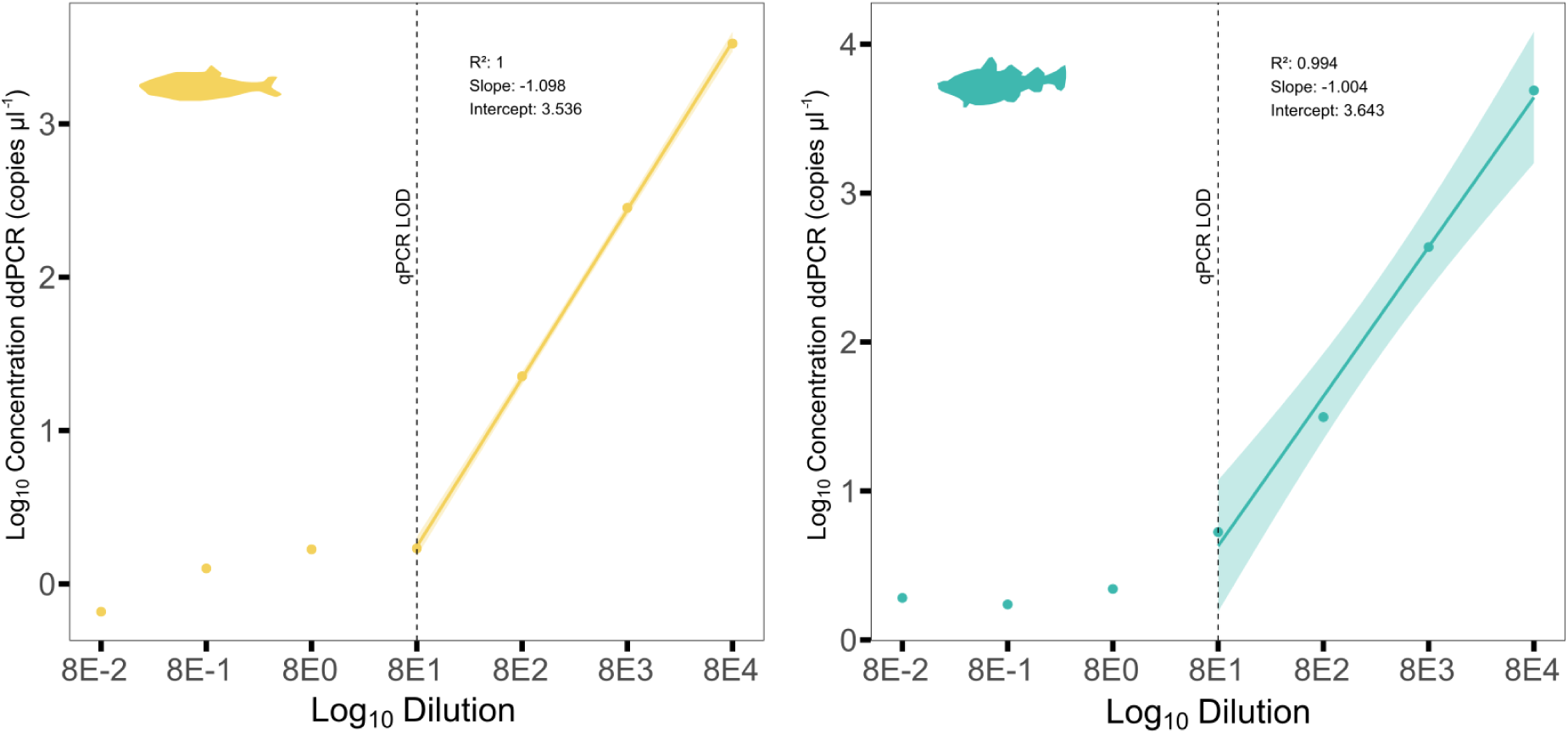
Standard curves generated from serial dilutions of tissue-derived DNA extracts for *Clupea harengus* (yellow) and *Gadus morhua* (turquoise), used to evaluate assay performance and detection sensitivity. Linear models were fitted to the four highest-concentration dilutions (indicated by solid lines), corresponding to the validated detection range based on qPCR limits of quantification (LOQ) established by Knudsen et al., (2019).

**Figure S3:**
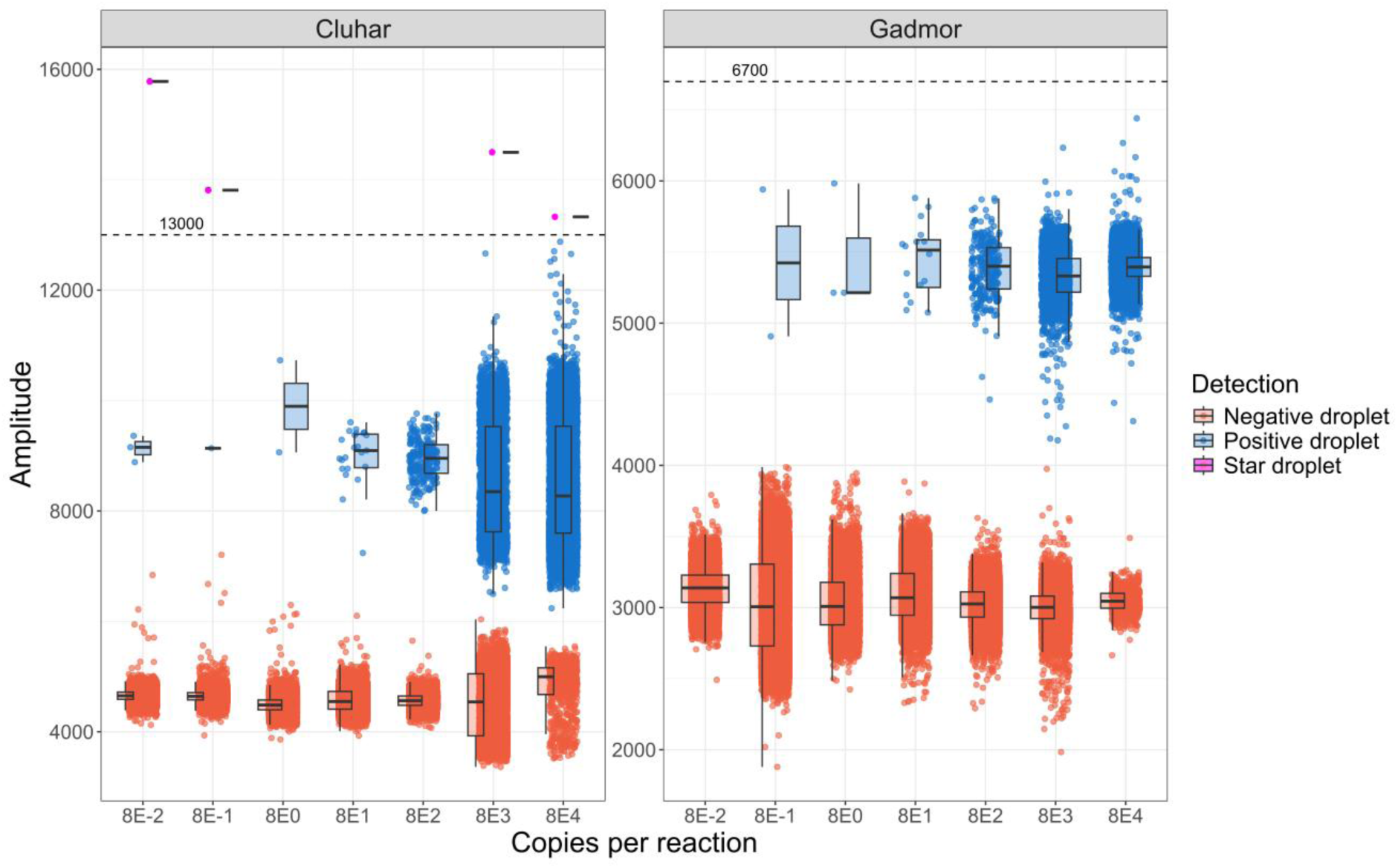
Fluorescence amplitude distributions across the standard dilution series for *Gadus morhua* and *Clupea harengus*. Positive (blue), negative (red), and intermediate (“star”) droplets (purple) are shown to illustrate threshold determination. Upper amplitude thresholds were conservatively set at 7,000 units for *G. morhua* and 13,000 units for *C. harengus* to exclude non-specific high-amplitude droplets.

**Figure S4:**
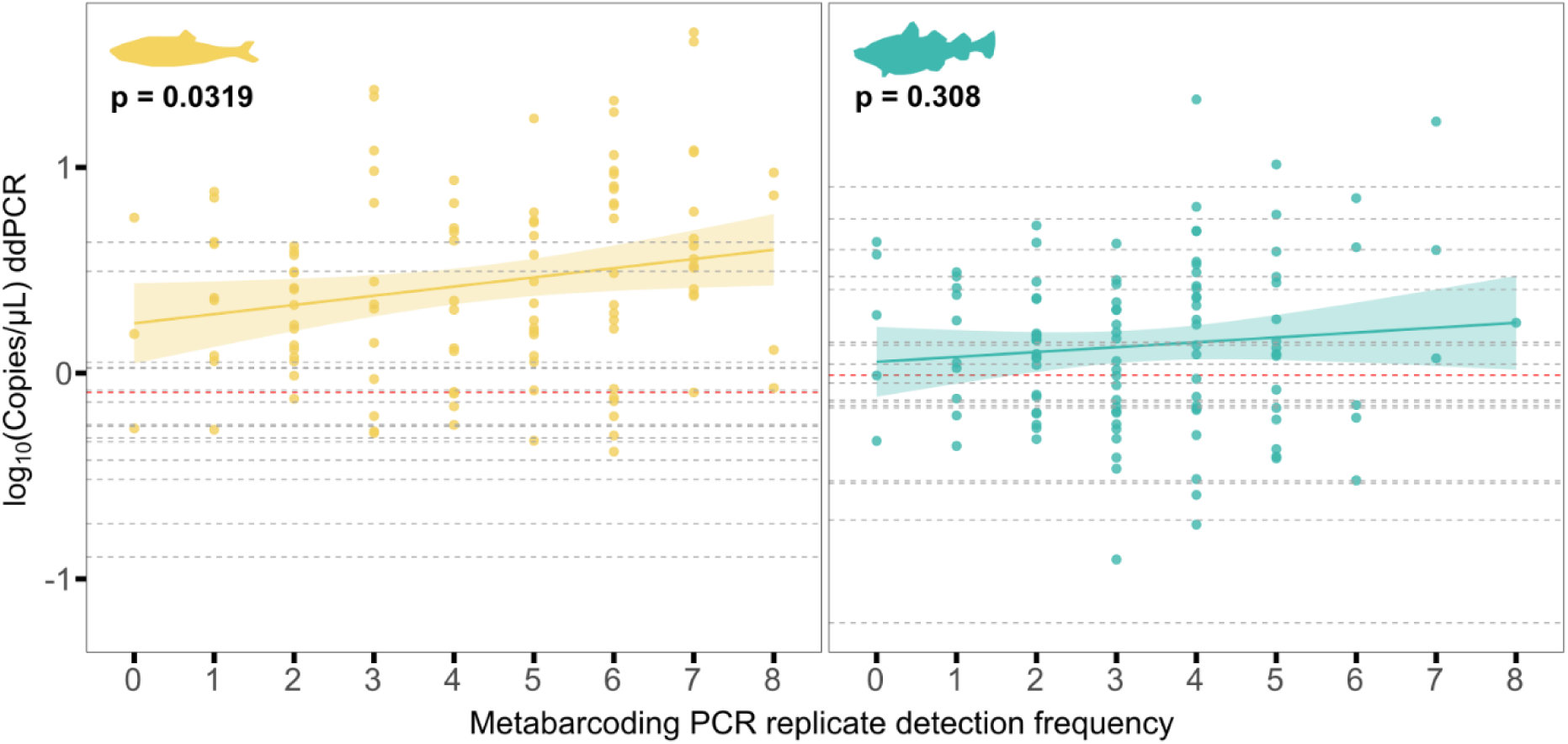
Relationship between ddPCR concentration and metabarcoding PCR replicate detection frequency for Atlantic herring (yellow) and Atlantic cod (turquoise) across sediment samples from two Icelandic marine sediment cores (PC019 and GC01; data combined). ddPCR concentrations (copies µL⁻¹) are shown on a log_10_ scale and plotted against the number of positive metabarcoding PCR replicates per sediment sample (out of eight technical PCR replicates). Grey points represent no-template controls (NTCs); red dashed lines indicate the species-specific mean ddPCR concentration measured in NTCs. Solid lines show linear regression fits with shaded 95% confidence intervals. Statistical significance (p-values) of each regression is indicated in the panels.

**Figure S5:**
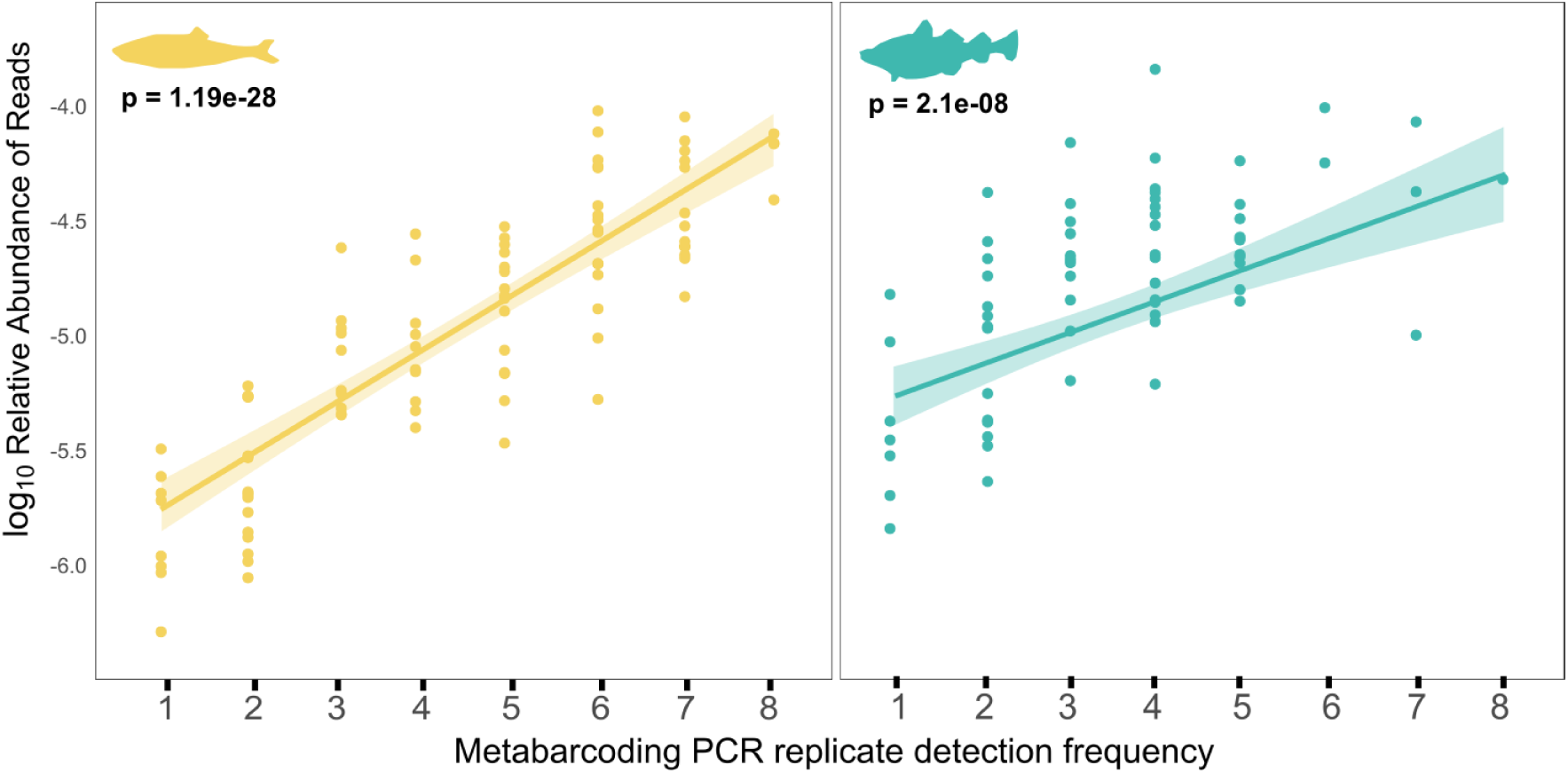
Proportion of positive metabarcoding PCR replicates versus log_10_-transformed relative read abundance for genus *Clupea* (yellow) and *Gadus* (turquoise). Statistical significance (p-values) of the correlations is indicated on each plot.

**Figure S6:**
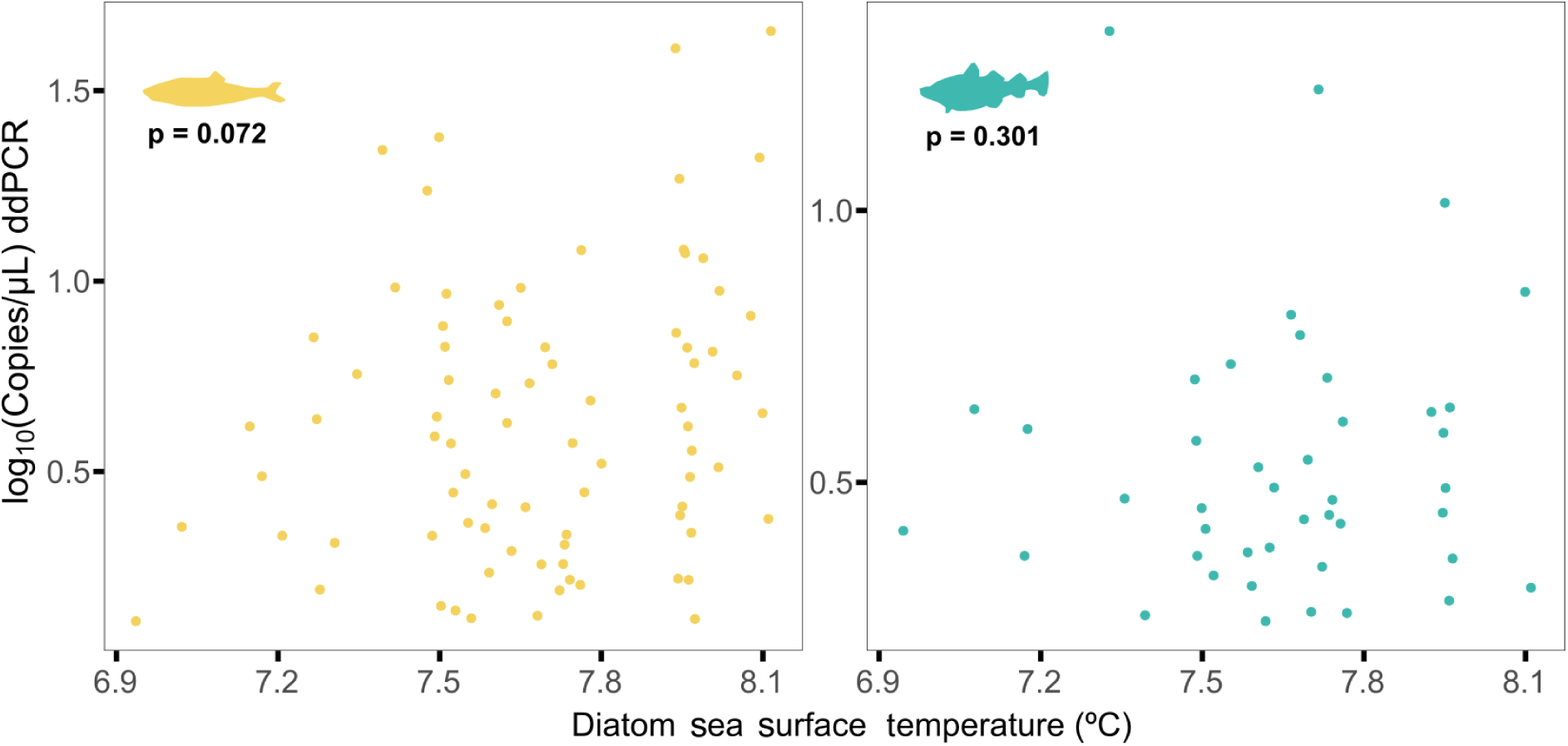
Log_10_-transformed ddPCR sedaDNA concentrations plotted against reconstructed sea surface temperature (SST) estimates derived from a diatom transfer function for Atlantic herring (*Clupea harengus*, yellow) and Atlantic cod (*Gadus morhua*, turquoise).

